# The gems of the Middle-East: Unveiling the biodiversity of Monogonont rotifers in temporary waterbodies of Israel

**DOI:** 10.1101/2023.10.25.563682

**Authors:** Ofir Hirshberg, Sofia Paraskevopoulou, Katrin Kiemel, Frida Ben-Ami

## Abstract

Temporary ponds represent ecologically important habitats that support high species diversity and provide essential ecosystem services, particularly in certain countries surrounding the Mediterranean Sea, where natural lakes are scarce. Israel is located along the southeastern Mediterranean coast and features Mediterranean and arid climatic zones that vary in a variety of meteorological parameters. Temporary ponds are prevalent throughout Israel, including the Mediterranean coast, Mediterranean mountain (i.e., Galilee region), and arid climatic zones. These temporal ponds harbor diverse invertebrate communities that exhibit significant spatial and temporal variations. Among these communities, Monogonont rotifers are notably one of the most diverse and abundant groups. Despite their significant role in aquatic food webs, rotifers are often overlooked in environmental studies, likely due to their small size and difficulties associated with their taxonomy. Resilient resting eggs produced by Monogonont rotifers during periods of unsuitable environmental conditions serve as significant source of propagules that drive the recolonization of temporary ponds upon rehydration, thereby influencing the dynamics of the pond community and metacommunity organization. Here, we examined the diversity of Monogonont rotifers by rehydrating sediment from 30 temporary ponds across Israel. Our analysis identified 39 species, with 25 (64%) of them being reported for the first time in Israel. We found the highest diversity of Monogonont rotifers in the Mediterranean coast region, which is characterized by low altitude, long hydroperiod, and relatively low mean summer daily maximum temperature, while the lowest diversity was found in the Arid region which is characterized by high altitude, short hydroperiod, and high mean summer daily maximum temperature. Our findings suggest that altitude, hydroperiod and mean summer daily maximum temperature are important parameters associated with the observed biodiversity patterns. Our metacommunity analysis further revealed a small contribution of geographic distance (2%) and environmental factors (1%) in shaping rotifer metacommunities. We also found a significant positive correlation among community composition, environmental distance (i.e., Gower’s distance) and geographic distance, possibly due to a linearity in the sampling set-up. Overall, our study highlights the importance of temporary ponds as significant habitats for diverse rotifer communities and emphasizes the need to further study “micro” invertebrate diversity in these unique ecosystems.

## Introduction

Ponds represent the most prevalent type of water bodies globally (Meerhoff & Jeppesen, 2009), albeit their ecological significance has only recently been recognized (Hill et al., 2021, Céréghino et al., 2014). These freshwater habitats, including temporary ponds known for their high species diversity and essential ecosystem services (Steward et al., 2017; Vico et al., 2020; Coutts et al., 2013), are among the most vulnerable habitats. They are particularly threatened by changes in hydrological patterns and the invasion of exotic species (Myers et al., 2000; Sala et al., 2000), making them highly susceptible to environmental changes and urbanization (Myers et al., 2000; Oerti et al., 2005; Sala et al., 2000).

The Mediterranean region, known for its temperate and arid climate, hosts a high density of temporary ponds which undergo periodic dryness (Zacharias & Zamparas, 2010). These temporary ponds harbor a diverse range of aquatic and semi-aquatic organisms (Bagella et al., 2016, Boix et al., 2016), comprising an important component of the Mediterranean aquatic ecosystems, especially in areas where natural lakes are scarce (Miracle et al., 2010).

Israel, positioned within the eastern Mediterranean basin, is characterized by Mediterranean and arid climate zones that vary mainly in their average temperature and precipitation (Fleischer et al., 2008). Temporary ponds are abundant throughout Israel, spanning from the temperate Upper Galilee in the north to the Mediterranean coastline and the desert in the south (Goren & Ben-Ami, 2013). In addition to its great habitat diversity (Cerdà, 1998), Israel is an important pathway for migratory birds (Frumkin et al., 1995), which are known to be an effective dispersal vector for some aquatic invertebrates (e.g., rotifers, crustaceans) (De Morais Junior et al., 2019; Simonis & Ellis, 2014; Green et al., 2008; Frisch et al., 2007). The appearance and potential dispersal of organisms carried by these dispersal vectors, coupled with the periodic occurrence of the temporary ponds, lead to alterations in both the landscape (e.g., resource links/resource allocation; Jeltsch et al., 2013) and the communities inhabiting these ponds (Altermatt & Ebert, 2010; Hirshberg & Ben-Ami, 2019). Besides these (external) biotic influences, abiotic factors such as hydroperiod length and habitat size may also affect the invertebrate biodiversity patterns observed in these habitats (Boix et al., 2016).

Rotifers (Phylum: Rotifera) are regularly found in temporary ponds and represent important links in aquatic food webs as primary consumers. They feed on unicellular organisms and serve as prey by other invertebrates and fish larvae (Brandl, 2005), thereby, transfering energy to higher trophic levels (Wallace, 2002). Rotifers comprise three subclasses, Seisonidea, Bdelloidea, and Monogononta (Segers, 2007), with the latter being the most species rich (> 1,500 species) and morphologically diverse group within the rotifer phylum (Segers, 2007).

Monogonont rotifers exhibit cyclical parthenogenesis, a reproductive pattern consisting of alternating phases of clonal reproduction interrupted by sexual reproduction when an environmental trigger is perceived. This sexual reproduction results to the formation of dormant stages called resting eggs (Gilbert, 2020). Among environmental conditions that trigger sexual reproduction, population density, diet, and photoperiod are the best documented (García-Roger et al., 2017; Gilbert, 2020). Resting eggs are very resilient and may remain in a viable dormant stage for a long time until favorable conditions are restored (Schröder, 2005). As a result, they accumulate in the sediment, forming the so-called “egg banks” and therefore representing a spatiotemporal biodiversity spectrum (Brendonck & De Meester, 2003). These characteristics allows these egg banks to serve as a protective measure against unfavorable conditions/extinction (Brendonck & De Meester, 2003; Wallace, 2002) and as a suitable dispersal agent for passive dispersal by wind or biotic vectors (i.e., mobile links) (Incagnone et al., 2015; Pinceel et al., 2016; Fonatento, 2019).

While historically rotifers were considered to have a ubiquitous biogeographic distribution (Fontaneto et al., 2012), recent studies have revealed environmental specializations even within evolutionary closely related species (e.g., Paraskevopoulou et al., 2018, 2020; Gabaldon et al., 2016; Zhang et al., 2019; Walczyńska & Serra, 2022). As certain rotifer species tolerate different levels of pollution and eutrophication, they are widely used as bioindicators (Dahms et al., 2011; Kuczynski, 1987; Wallace, 2002). Beside their ecological importance, rotifers are of high economic value in aquaculture, providing a high-quality food source for fish larvae (Garcia et al., 2008; Hagiwara & Yoshinaga, 2017; Lubzens et al., 1989). In Israel, Monogonont rotifers are extensively used in mariculture industry (Lupzens et al., 1997). Additionally, their biodiversity is being monitored in lake Kinneret, the only permanent large waterbody in Israel, by the Israel Oceanographic and Limnological Research (IOLR). Nevertheless, little is known about the diversity, species richness and abundance of monogonont rotifers in the vast majority of Israel’s temporary ponds.

To understand the potential impact of forthcoming environmental changes on temporary ponds and their primary consumers, it is crucial not only to assess their biodiversity but also to understand the underlying environmental factors shaping the observed biodiversity patterns (Gontier et al., 2006). Metacommunity ecology is a scientific field that attempts to link local biodiversity patterns to the patterns we observe at a regional scale. The underlying dynamics are embodied in metacommunity theory, which was first introduced by Leibold et al. (2004) and has since then further developed (Leibold & Chase, 2018). This theory aims to identify the processes governing metacommunity dynamics, i.e., dispersal limitation, environmental filtering/niche selection, species interaction and stochasticity/ecological drift (Leibold & Chase, 2018; Thompson et al., 2020; Vellend, 2010), to derive predictions on communities under anthropogenic/environmental change (e.g., climate change, pollution, habitat fragmentation; Chase et al., 2020). While the metacommunity theory has been applied to various freshwater systems (e.g., river systems; Ramos et al., 2022; Zhao et al., 2017, pond systems; Gálvez et al., 2020; Cottenie et al., 2003; Kiemel et al., 2022, rock pools; Vanschoenwinkel et al., 2010) and geographical areas (e.g., Gálvez et al., 2020; Cottenie et al., 2003; Kulkarni et al., 2019), temporary ponds in Israel have not yet been analyzed in the context of the metacommunity framework. Understanding the underlying processes could be of great importance in establishing future management strategies, as pond habitats of Israel may shrink due to changes in land use, urbanization, and climate change (Levin et al., 2009).

This study attempts to address this knowledge gap by analyzing the communities of monogonont rotifers in 30 temporary water bodies in Israel by rehydrating resting eggs stored in sediments. Our aim is (I) to determine the overall biodiversity of monogonont rotifers in Israel, (II) to identify key environmental drivers shaping abundance and biodiversity, and thus (III) to draw conclusions on the underlying processes that shape the monogonont rotifer communities across different climatic zones.

## Material and Methods

### Study area

The study area is a 15,600 km^2^ area of Israel that comprises three main climatic regions (*Figure 1*). Each of the different regions harbors different climate and topography. The upper Galilee region has a Mediterranean climate and mountain topography, hereafter referred to as the ‘Mediterranean mountain’ region, while the Mediterranean coastal plain also experiences a Mediterranean climate, but it is located on the Mediterranean coast, and it will thus be referred to as the ‘Mediterranean coast’ region. The Negev Desert will be referred to as the ‘Arid’ region, because it is dominated by an arid/desert climate. The ponds located in the Mediterranean coast and Mediterranean mountain regions are usually large (375-200,000 m^2^) and maintain water for most of the year, with many of them located in urban nature reserves or parks. Some of these ponds are located in proximity to one another and may connect in high precipitation seasons. Ponds located in the arid region are highly ephemeral and may desiccate several times a year.

**Figure 1:**
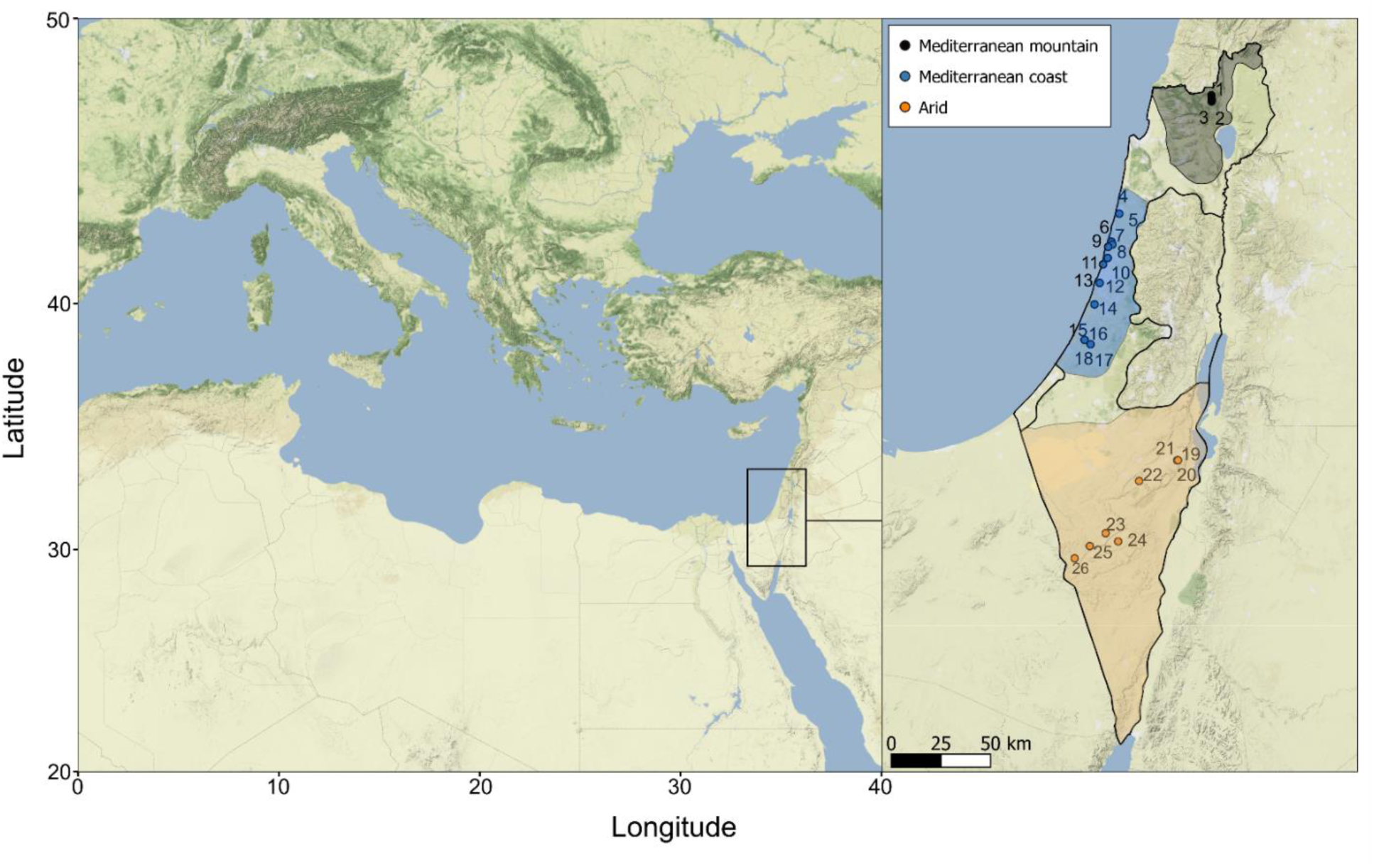
Map of the sampled water bodies across three climatic zones of Israel. Sampling locations within the three climatic zones, Mediterranean mountain (black), Mediterranean coast (blue), and Arid (orange).

### Sampling

During the dry period of 2019 (August-November), sediment samples were collected by scratching of the surface (i.e., the first 5 cm are usually considered as the active egg bank; Brendonck & De Meester, 2003) of various locations in the pelagic and littoral areas of 30 water bodies in different regions of Israel (Mediterranean coast, Mediterranean mountain, and Arid; *Figure 1*). In total we sampled approximately 500 g of sediments, which were packed in zip-lock plastic bags, labeled, and stored at room temperature (20 ^°^C) in the dark until further processing.

### Hatching experiment and species identification

For the sediment rehydration and hatching experiment, sediment from each site was homogenized by mixing it thoroughly. Then, 20 g of the sediment were placed in a 300 mL jar with 250 mL of either WC medium (Guillard & Lorenzen 1972) or artificial *Daphnia* medium (ADaM, Klüttgen et al., 1994; Ebert et al., 1998). One replicate for each hatching medium was processed to account for potential hatching biases introduced through the hatching medium. The resulting 60 jars were placed in a climate chamber (Snijders ECP01E, Netherlands) at 20 ^°^C and a 12:12 h L:D cycle while humidity was set to 60 %. After 7 days, the samples were filtered using a 40 µM mesh size filter and screened for rotifers under a dissecting microscope (Leica 205C, Germany). Screening was performed once per week for a total period of four weeks. This threshold was set as preliminary experiments showed that prolonging the experiment beyond four weeks did not yield a significantly higher number of species that hatched from resting eggs.

Morphological species identification was performed by using a phase contrast microscope (Leica 2500, Germany), and a combination of several identification keys (Fischer and Ahlrichs, 2011; Koste, 1978; Nogrady et al., 1995; Nogrady and Segers, 2002; Segers, 1995; Segers and Walsh, 2017; Wallace et al., 2019) and catalogs (Jersabekand & Leitner, 2013). When information regarding the size of the rotifers and their different body parts was required for a successful species level identification, measurements were collected using a microscope camera (Leica DFC 295, Germany). To determine morphological identification at the species level, in genera such as *Cephalodella,* where a closer observation of the trophi is required, rotifers were placed in 88% lactic acid (CH_3_CH(OH)COOH) to dissolve the soft tissue (örstan, 2015).

### Statistical analysis

All statistical analyses were performed using R version 4.1.1 (R Core Team, 2020). To analyze hatched species compositions in regard to spatial and environmental parameters (*Supplementary material table S1*), daily/annual precipitation, mean summer daily maximum temperature, mean winter daily minimum temperature, near ground temperature, daily/annual evaporation and wind velocity were retrieved from the Israeli Meteorological service (https://ims.gov.il). The size of the ponds (m^2^) was acquired from previous surveys (Elron & Gafni, 2011) or estimated in the field. Longitude, latitude, altitude, and number of neighboring ponds were obtained by extracting GPS data and using Google Earth. To obtain a reliable proxy for hydroperiod (i.e., permanency) of water containment, ratios of evaporation/precipitation (e/p) and size/evaporation/precipitation (s/e/p) were calculated (LaBaugh et al., 2016). For some summary statistics (i.e., ANOVA, NMDS), permanency and altitude were grouped into five categories (details see *Supplementary material table S2*).

### Species co-occurrence and diversity indices

To assess potential functional co-occurrence patterns the *cooccur* function of the package cooccur v1.3 (Griffith et al., 2016) was used. Species richness and species diversity was examined using three indexes, which differentially emphasize species richness, abundance, and evenness (i.e., species richness, Shannon index, and Simpson index). The diversity indices were calculated using the *spec*.*number* and *diversity* functions of the package vegan 2.6-4 (Oksanen et al., 2013). To identify potential relationships between environmental parameters and diversity indices, parameters (i.e., size, altitude, number of neighboring ponds, permanency (s/e/p; details see *Supplementary material table S1*), mean summer/winter daily maximum/minimum temperatures respectively, and wind velocity) were analyzed using a generalized linear model (GLM). To avoid overfitting, correlation between the explanatory variables was examined using Pearson’s correlation coefficients with a threshold of > 0.7. Non-correlated variables were then analyzed using the *glm.nb* function in package lme4 version 1.1 (Bates et al., 2023) for species richness and the *glm* function in package glmmADMB version 0.8.3.3 (Bolker et al., 2012) for Shannon and Simpson indices. In order to obtain a distribution similar to normal, the Shannon and Simpson indices as well as response variables were Tukey transformed before running the GLM. To identify the model that better explained the variance, model selection was based on the corrected Akaike information criterion (cAIC) utilizing the function *dredge* from the R package MuMIn (Barton, 2016). Normal distribution of model parameters was subsequently checked using the DHARMa package version 0.4.6 (Hartig, 2017).

### Community structure and metacommunity analysis

In order to assess the differences in community structure between temporary ponds from different regions differing in altitude, temperature, and permanency, we used a nonmetric multidimensional scaling (NMDS) approach based on the Jaccard (for presence/absence data) dissimilarity in combination with a Pemanova which was conducted based on 999 permutations. We used the presence/absence data in order to account for potential biases in abundance introduced by our hatching parameters (e.g., monitoring and counting abundance once per week). Analysis was performed using the *metaMDS* function of vegan package version 2.6-4 and the *pairwise.perm.manova* function on the RVAideMemoire 0.9-81-2 package (Hervé & Hervé, 2020).

To identify potential effects of environmental parameters on the community composition of the temporary ponds, a distance-based redundancy analysis (dbRDA) was used based on the Jaccard dissimilarity, using the *dbrda* function in vegan version 2.6-4 package (Oksanen et al. 2013). To avoid overfitting, correlation coefficients were calculated using Pearson’s correlation coefficients with a threshold > 0.7. Non-correlated explanatory variables (i.e., size, altitude categories, mean summer daily maximum temperature, wind velocity, permanency (s/e/p), neighboring ponds at 1 km, 5 km and 15 km radius) were subsequently fed into a forward stepwise selection using 9999 permutations implemented in the *ordistep* function in vegan v.2.6-4 package (Oksanen et al., 2013). In order to include distance between the ponds as an explanatory variable, principal coordinates of neighboring matrices (PCNM) were calculated based on the coordinates of the different ponds, using the *pcnm* function in vegan v.2.6-4 package (Oksanen et al., 2013). These PCNMs were then fed into a forward selection dbRDA to identify significant PCNMs. Resulting significant environmental (i.e., mean summer daily maximum temperature, altitude, and permanency (s/e/p)) and spatial parameters (i.e., PCNM1 and PCNM2) underwent a variance partitioning (i.e., a method often used to detect underlying metacommunity dynamics; Bocard et al., 1992, Cottenie, 2003; Zhao et al., 2017, Kiemel et al., 2022) using the *varpart* function in vegan v.2.6-4 package (Oksanen et al., 2013). Variance proportions were further tested for significance using an ANOVA-like permutation test (*anova.cca* function) with 999 permutations. To further correlate community dissimilarities (based on Jaccard dissimilarities) with geographic distance, we performed a Mantel test with 9999 permutations using the function *mantel* in vegan v.2.6-4 package (Oksanen et al., 2013). Additionally, the community dissimilarities were correlated using a Mantel test with a proxy of environmental parameters calculated on all available parameters (i.e., size, altitude categories, mean summer daily maximum temperature, wind velocity, permanency (s/e/p), neighboring ponds at 1 km, 5 km and 15 km radius) as the Gower distance (Gower, 1971).

## Results

### Species composition

We identified 39 monogonont rotifer species, comprising 20 genera, which were found in 26 out of the 30 investigated temporary ponds in Israel (*Table 1*), covering 10 families of the order Ploima and three of the order Flosculariaceae. Twenty-five out of the 39 identified rotifer species in this survey (64%) are reported for the first time in Israel. The family Brachionidae (Ploima) included nine species, six of which were of the genus *Brachionus*, distributed across all regions. The family Notommatidae (Ploima) included nine species, all of which were of the genus *Cephalodella*, and were also found in all regions. They were followed by the family Lecanidae (Ploima), which included a total of five species of the genus *Lecane*, exclusively found in the Mediterranean coast region. All other families were represented by 1-2 species with the majority of the species noted in the Mediterranean coast region (*Table 1*). The most common species was *Filinia longiseta* (Flosculariaceae: Trochosphaeridae) which was present in 11 ponds, dominantly in the Mediterranean coast region. *Filinia longiseta* was followed by the species *Hexarthra mira* (Flosculariaceae: Hexarthridae) which was recorded in the Mediterranean coast and arid regions of Israel for the first time, and *Epiphanes brachionus* (Ploima: Epiphanidae), which was found in the Mediterranean coast and Mediterranean mountain regions. The species *Polyarthra vulagris* (Ploima: Synchaetidae) and *Brachionus quadridentatus* (Ploima: Brachionidae) were also common, mainly in the Mediterranean coast region, being present in nine ponds. *Brachionus quadridentatus* together with *Brachionus varaibilis* were also reported for the first time from Israel. A few smaller sized species of the genera *Cephalodella*, *Lepadella*, and *Trichocerca* were also reported for the first time from the region (*Table 1*). Importantly the species *Rhynoglena ovigera* (Ploima: Epiphanidae) reported in two out of the three ponds in the Mediterranean mountain region represents the first record of the species outside of the Nearctics.

**Table 1:**
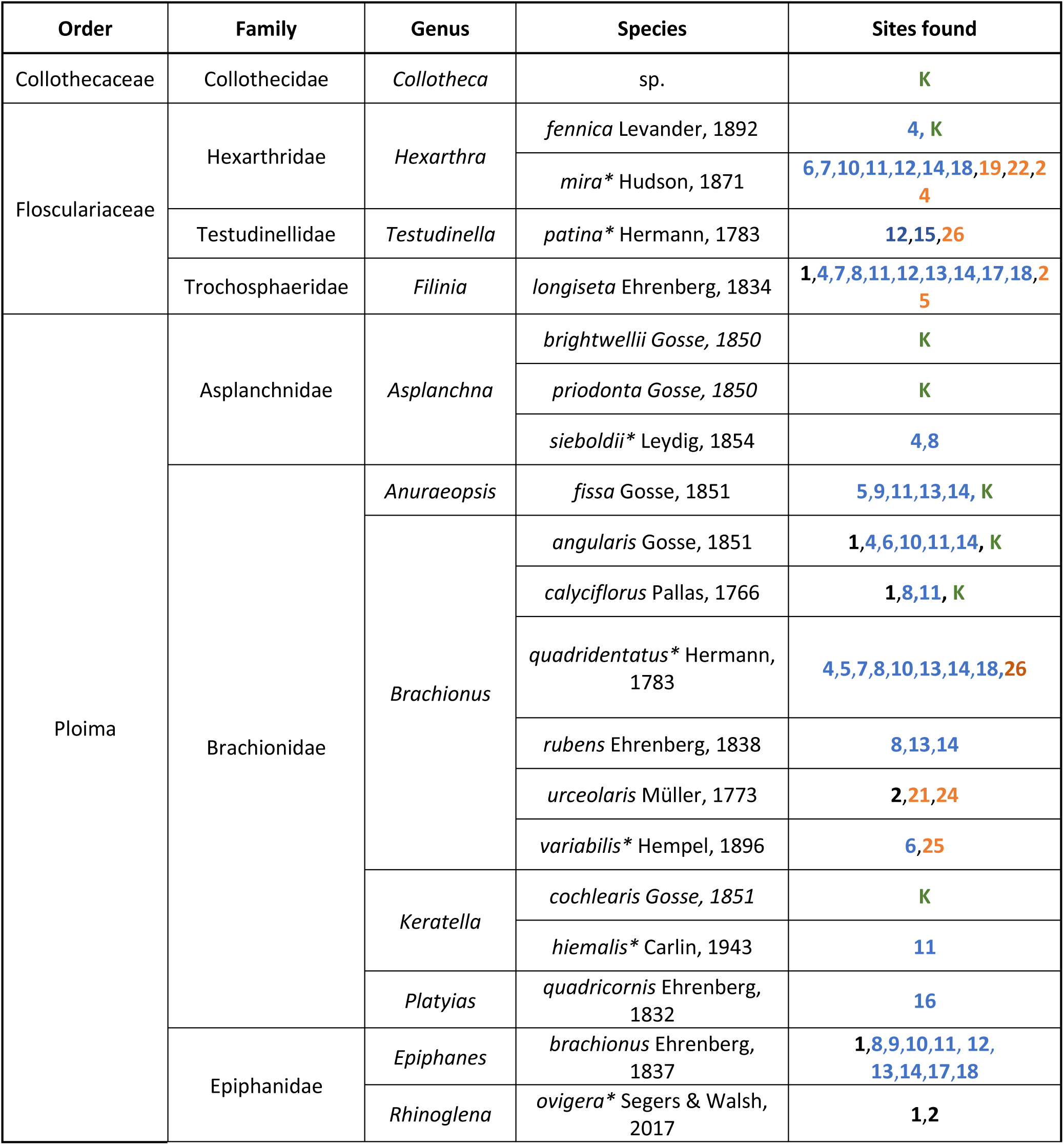
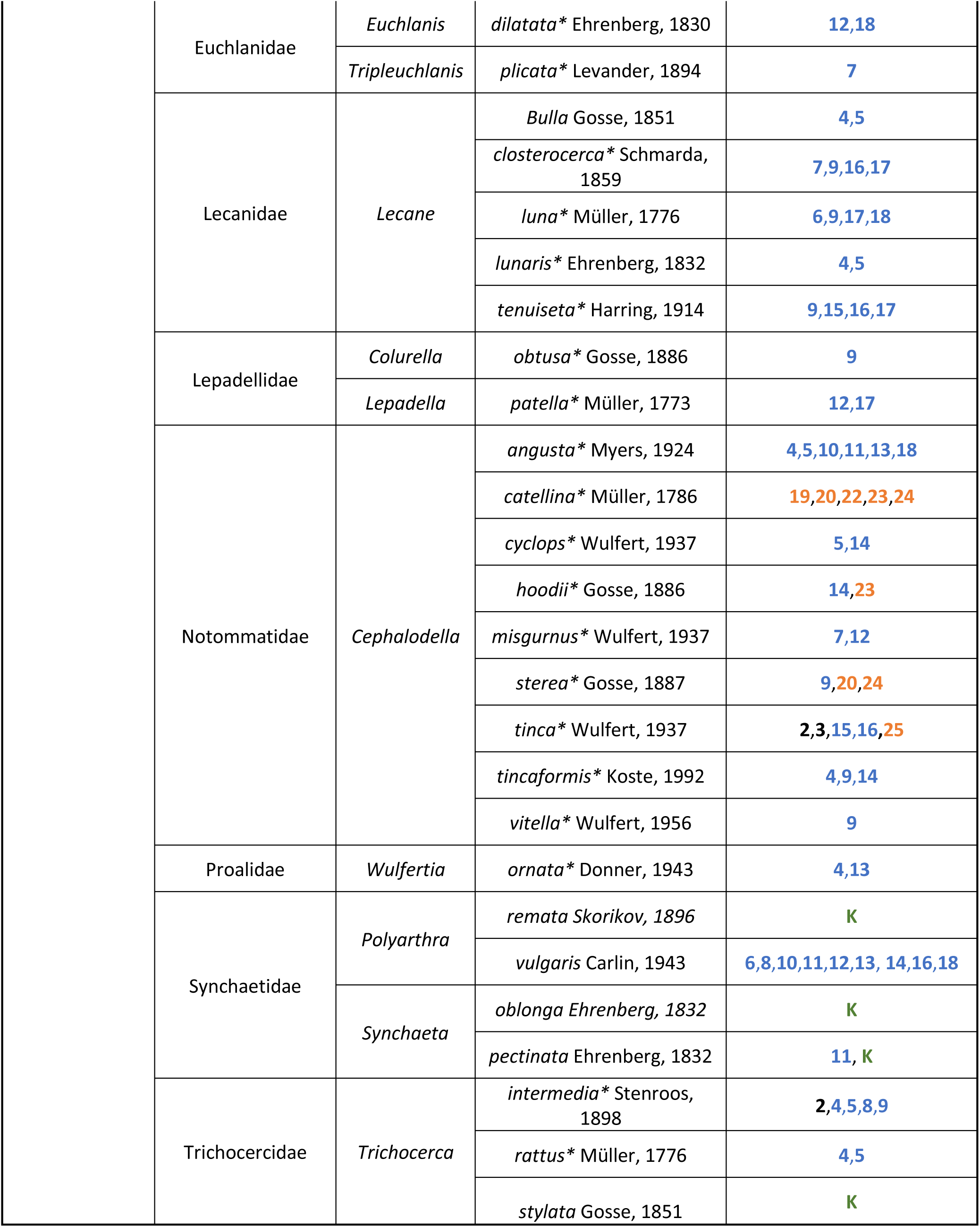
List of Monogonont rotifer species recorded in Israel. Asterisks indicate species that are reported for the first time from Israel during our survey. Number abbreviations correspond to ponds depicted in *Figure 1*. Species found in Lake Kinneret (K) were taken from past publications (Gal & Hambright, 2014; Gal et al., 2020; Gophen, 2005). *Brachionus calyciflorus* specimens were identified to the species complex level.

### Species co-occurrence

Co-occurrence patterns revealed that most species co-occurred randomly (*Figure 2*). However, two pairs of species showed a negative co-occurrence, *F. longiseta* and *Cephalodella catellina* (Ploima: Notommatidae) as well as *E. brachionus* and *Cephalodella tinca* (*Figure 2*). Positive co-occurrence patterns were detected in eight pairs of species (*Figure 2*). *Epiphanes brachionus* positively co-occurred with four species, *P. vulgaris, Brachionous calyciflorus, F. longiseta,* and *Brachionus rubens*, while *B. rubens* co-occurred additionally with *P. vulgaris* and *B. quadridentatus*.

**Figure 2:**
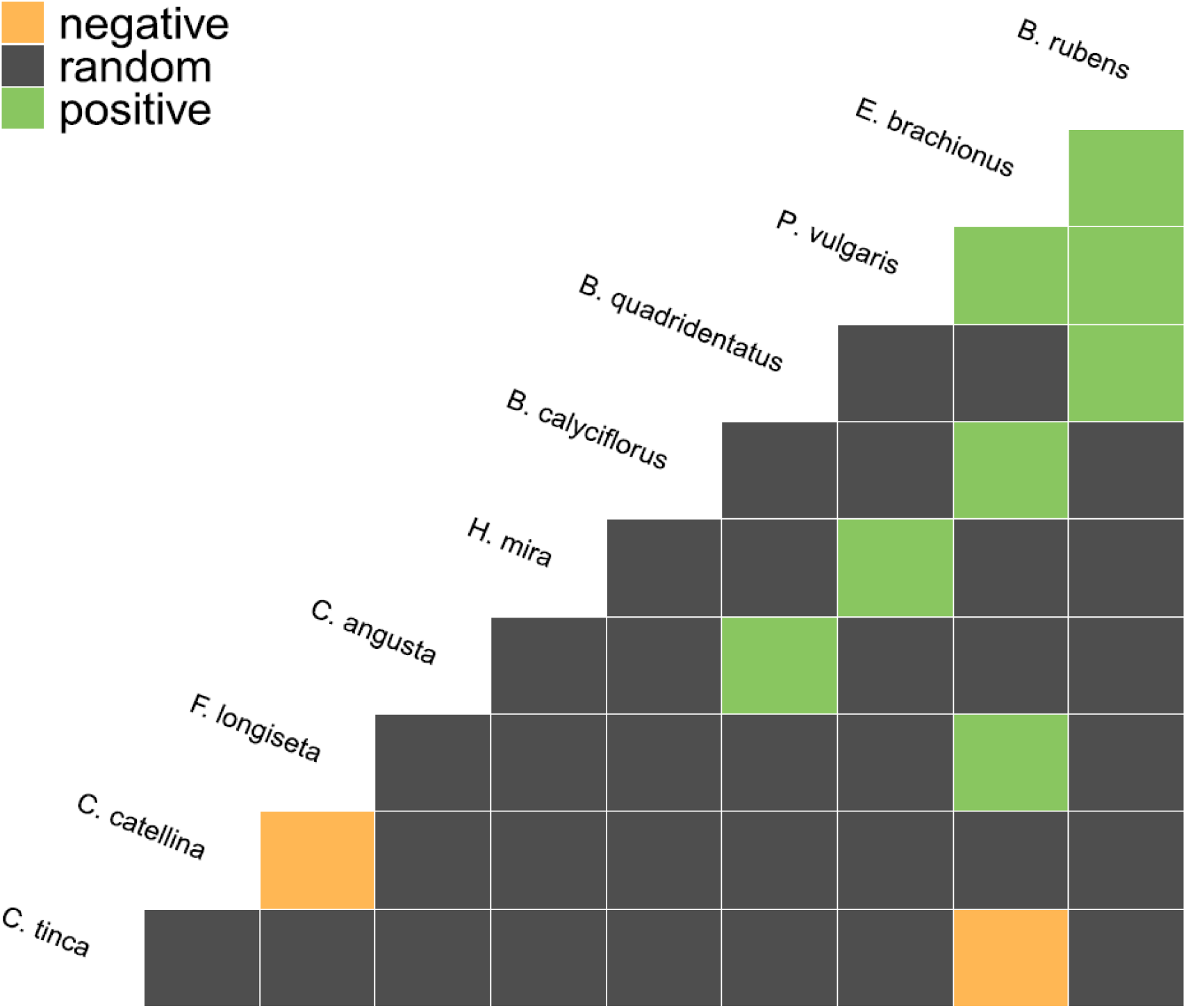
Calculated co-occurrence patterns of identified rotifer species within temporal ponds. Green color indicates positive co-occurrence, orange color indicates negative co-occurrence and gray color indicates a random co-occurrence of species. Only pairs of species with positive or negative co-occurrence are shown.

### Species diversity

Species richness was significantly higher in the Mediterranean coast region than in the Mediterranean mountain and arid regions (Wilcoxon: Mediterranean mountain: p = 0.02, arid: p < 0.001; *Figure 3a*). The GLM revealed a negative effect of mean summer daily maximum temperature and altitude on species richness (*Figure 4a, 4b*). Both the Shannon and Simpson diversity indices were significantly higher in the Mediterranean coast region than in the arid region (Wilcoxon: Shannon: p = 0.004, Simpson: p = 0.02; *Figure 3b, 3c*) but did not differ from the Mediterranean mountain region (Wilcoxon: Shannon: p = 0.3, Simpson: p = 0.57; *Figure 3b, 3c*). Similar to species richness, the Shannon and Simpson diversity indices were negatively affected by mean summer daily maximum temperature (*Figure 4c*, *4d*).

**Figure 3:**
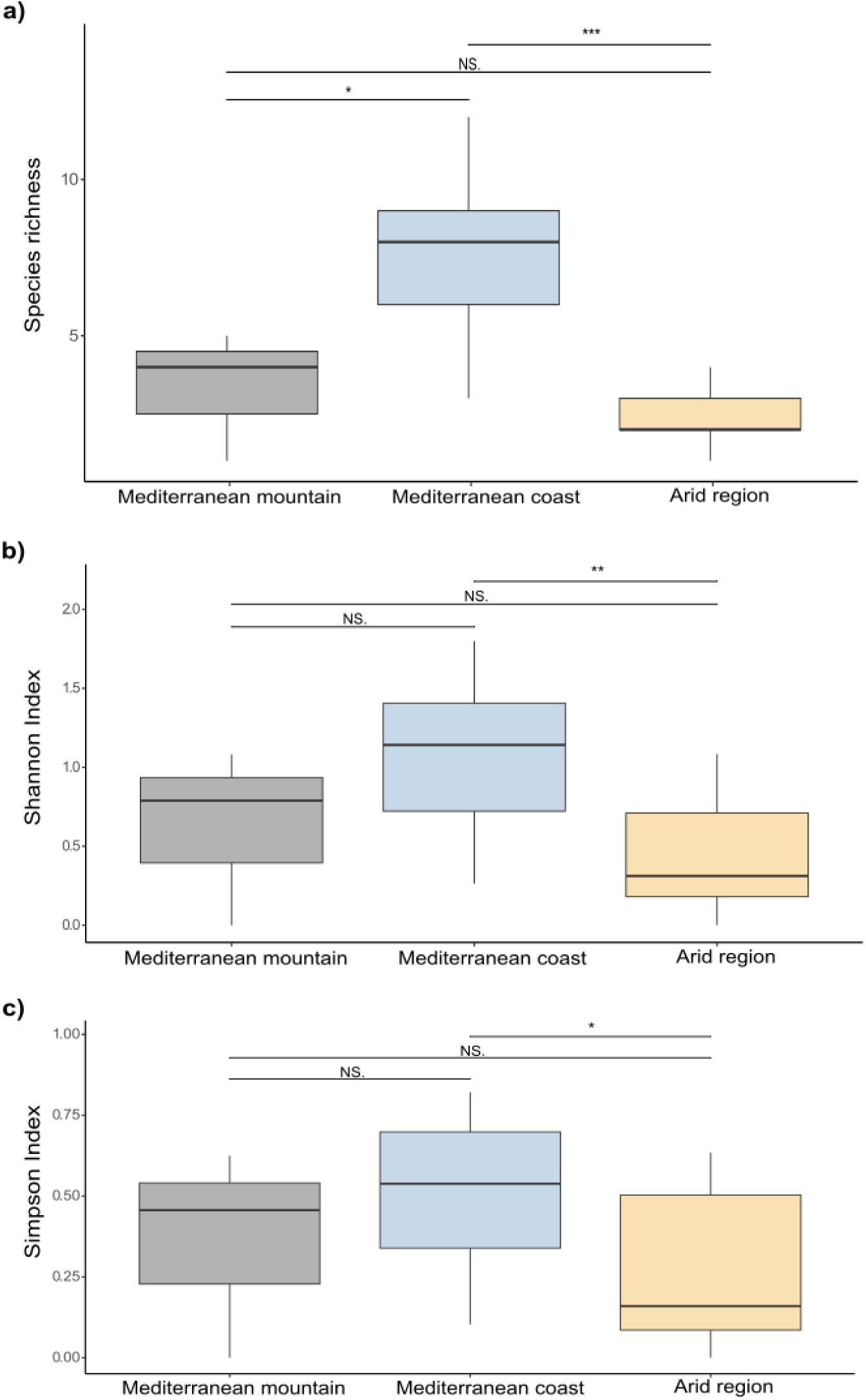
Comparisons of different diversity indices among the three study regions. A) species richness, B) Shannon index, C) Simpson index. Black, blue and yellow colors indicate Mediterranean mountain, Mediterranean coast and arid regions, respectively. Significance of pairwise Wilcoxon test is represented by brackets and asterisks (*; p<0.05, **; p<0.01, ***; p<0.001). Non-significant comparisons are indicated by NS.

**Figure 4:**
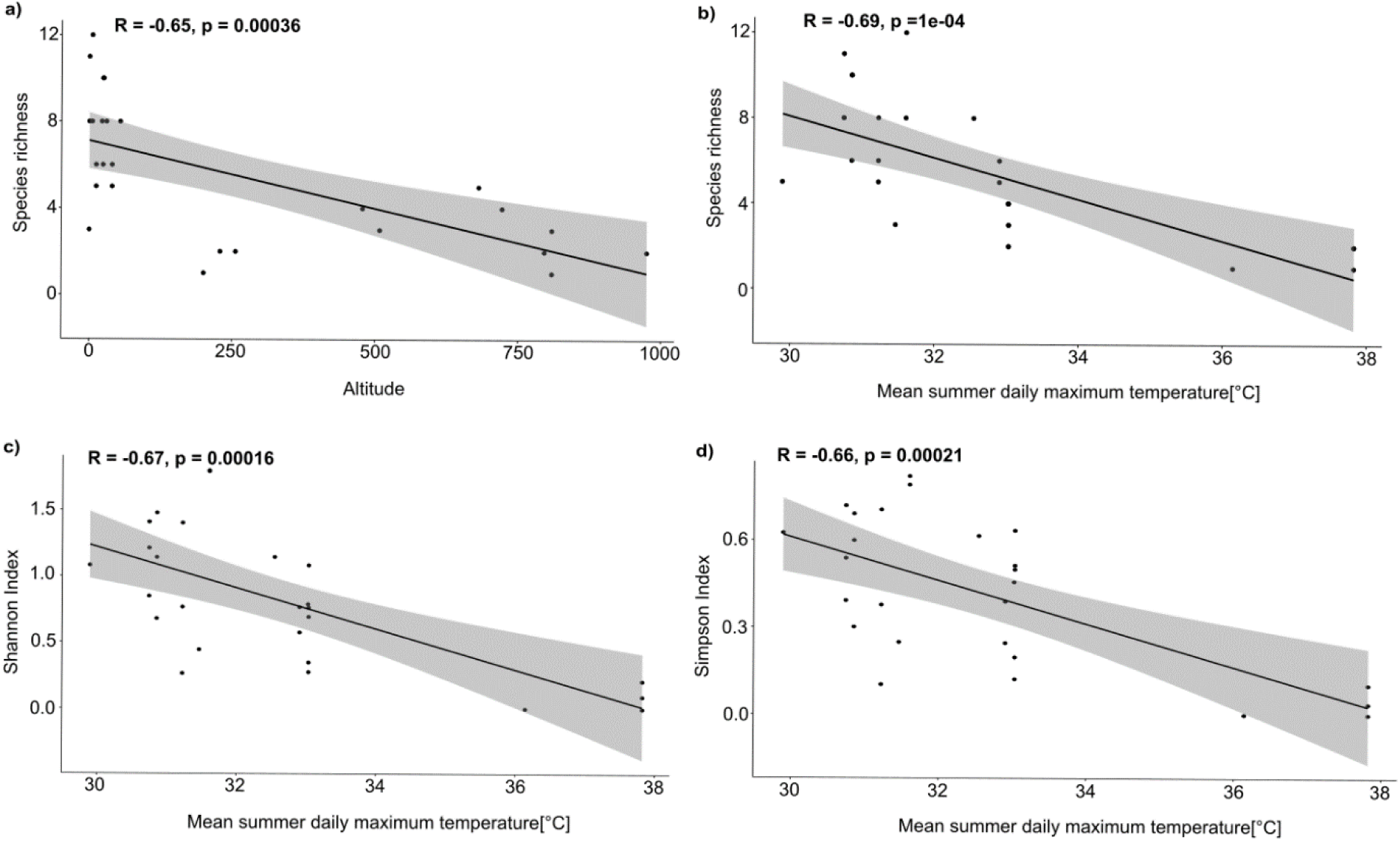
Generalized linear model showing the effects of environmental parameters on estimated diversity indices: a) altitude effect on species richness, b) mean summer daily maximum temperature effect on species richness, c) mean summer daily maximum temperature effect on Shannon index, c) mean summer daily maximum temperature effect on Simpson index.

### Community composition

The comparison of the community composition based on the presence absence data (i.e., Jaccard dissimilarity) revealed significant differences in the rotifer communities between the Mediterranean coast and mountain regions (p = 0.042) as well as between the Mediterranean coast and arid regions (p = 0.003). Rotifer communities were further influenced by altitude and permanency (s/e/p) (PERMANOVA: altitude: p = 0.06, permanency: p = 0.02; *Figure 5*). The dbRDA analysis of the rotifer communities based on present absence data (i.e., Jaccard dissimilarities) revealed that the communities were affected by mean summer daily maximum temperature (ANOVA: F = 2.77, p = 0.005) as well as the altitude (ANOVA: F = 1.61, p = 0. 05) and permanency based on s/e/p (ANOVA: F = 1.5, p = 0.045). Subsequent variance partitioning, based on significant spatial parameters (i.e., PCNM1 and PCNM2) and environmental parameters (i.e., mean summer daily maximum temperature and altitude categories), revealed an overall low explained variance by both, spatial and environmental parameters. Spatial components accounted for a higher proportion of variance (2 %) than environmental factors (1 %), leaving a large proportion of unexplained variance (92 %) (*Figure 6*). While the variance partitioning did not yield clear effects on environmental and spatial parameters, the subsequently conducted mantel test based on geographic distance and environmental distance (i.e., Gower’s distance), revealed a weak but significant positive correlation between community composition and geographic distance (Jaccard: R=0.119, *p*=0.03), as well as between community composition and environment (Jaccard: R=0.161, *p*=0.007) (details see *Supplementary material Figure S1, S2*).

**Figure 5:**
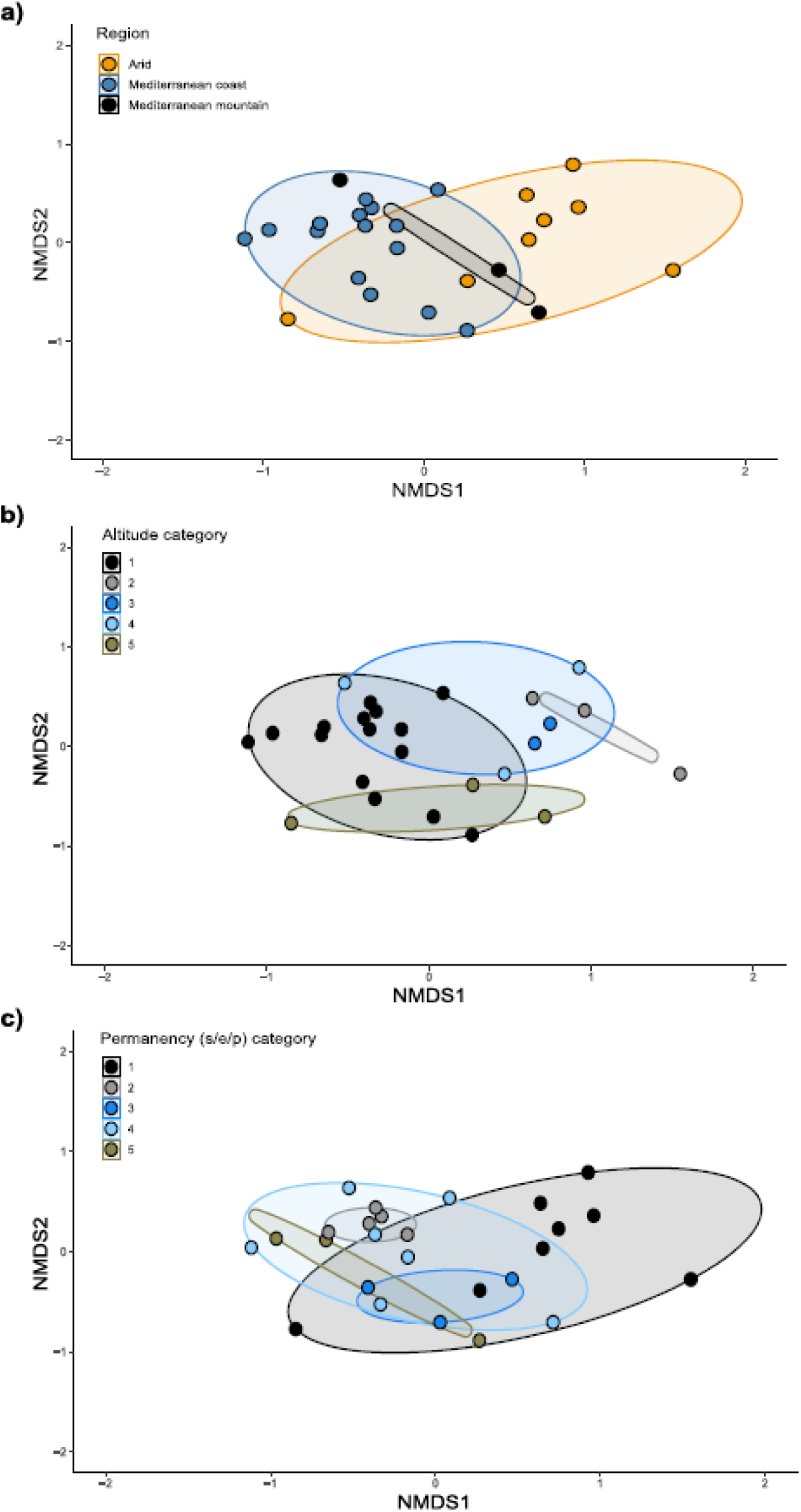
NMDS ordination plots representing the differences in community compositions based on Jaccard dissimilarities among the sampled temporary ponds: (a) regions, (b) altitude, and (c) permanency categories. Stress value=0.16.

**Figure 6:**
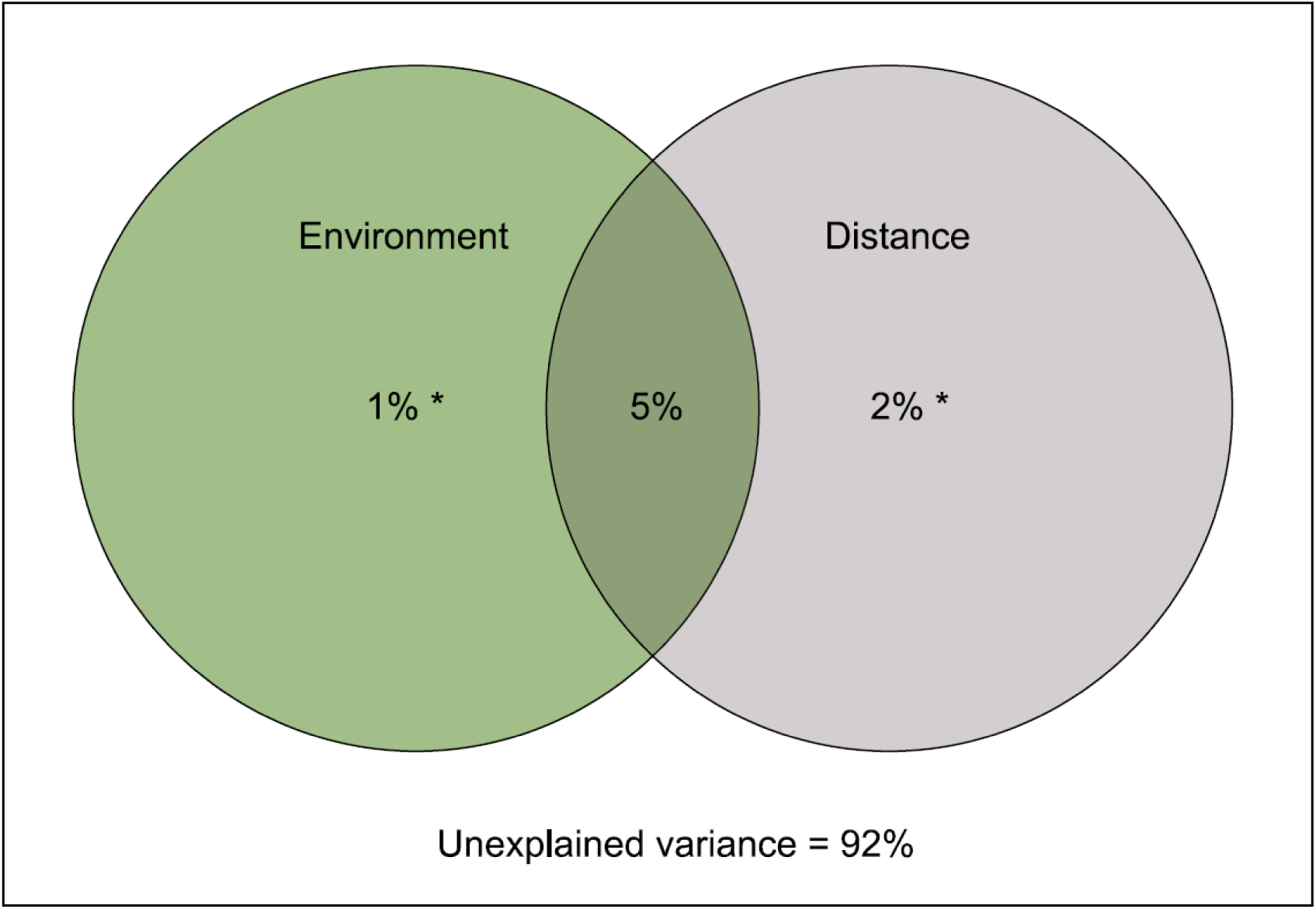
Variance partitioning based on significant parameters revealed by a dbRDA on the Jaccard dissimilarity between the ponds. Environment contains the parameters mean summer daily maximum temperature and altitude and distance is based on significant PCNMs (PCNM1 and PCNM2). Asterisks indicate significant fractions: * ≤ 0.05, ** ≤ 0.01, *** ≤ 0.001.

## Discussion

Our study demonstrates that Israel’s temporary ponds offer an ideal study area and present an opportunity for investigating how environmental factors affect the distribution of rotifer species within a small geographical area boasting diverse climatic zones. Overall, we found a total number of 39 monogonont rotifer species belonging to 20 genera. Although our study detected a lower number of rotifer species compared to other large-scale studies conducted in Italy, Russia, Poland, Mexico and USA (Braioni & Gelmini, 1983; Kutikova, 1998; Ejsmont-Karabin et al., 2004; Sarma et al., 2021; Brown et al., 2020), our findings suggest that a single sampling of spatio-temporal reservoirs of resting eggs can detect a sufficiently high number of species for diversity assessment and related community/metacommunity analysis. Most importantly, many of the species we detected (25) have not been previously reported from the region, substantially adding to the rotifer diversity in Israel. To date, the majority of recorded instances of rotifers in Israel originate from Lake Kinneret (Gophen, 2005; Gal & Hambright, 2014), which is the only large permanent waterbody in the country. Despite both being freshwater ecosystems, the composition of rotifer species in temporary ponds is notably different from that of Lake Kinneret. Lake Kinneret predominantly hosts planktonic rotifers (Gophen, 2005) whereas in temporary ponds we also found non-planktonic rotifer genera such as *Cephalodella* and *Lecane*. These disparities in species composition can be attributed to the distinct sampling methods. Sampling in Lake Kinneret involves using nets at various depths in the water column, primarily targeting exclusively pelagic organisms (Gal et al., 2020). Conversely, our study involved the rehydration of resting eggs that lay on the sediment, enabling us to capture a wider range of rotifer species, including planktonic, benthic, and occasionally periphytic species. Another possible explanation for the differences in detected species might be the habitat differences such as the presence of predators in a system. While Lake Kinneret is inhabited by planktivorous fish, such as *Mirogrex sp*. and *Acanthobrama sp*., which may prey on rotifers (Gophen & Serruya, 1990), temporary ponds are mostly fishless.

Among the most highly represented species in the 26 temporary ponds were *Hexarthra mira*, *Polyarthra vulgaris* and *Brachionus quandridentatus*. These species comprise species-complexes (García-Morales & Domínguez-Domínguez, 2019; Moreno et al., 2017; Zhang et al., 2021) which suggests that their widespread distribution among different climatic regions may be attributed to the presence of different cryptic entities with distinct ecological requirements. This phenomenon has been previously observed in other rotifer taxa such as *Brachionus plicatilis* and *Brachionus calyciflorus*, which were once thought to have a ubiquitous distribution, but were later found to be species complexes with relatively restricted ecological niches (Gabaldón et al., 2017; Zhang et al., 2018; Walczyńska et al., 2023). Apart from the Mediterranean coast region, the species *H. mira* was prevalent in most ponds within the arid region. Rotifers of the genus *Hexarthra* are typical inhabitants of ephemeral ponds in other geographical areas (i.e., Texas, USA; Schröder et al., 2007) and are characterized by a rapid growth and resting egg development (Schröder et al., 2007). These traits likely allow *H. mira* to thrive in the challenging and unpredictable environments of temporary ponds. Another common inhabitant of the arid region was species of the genus *Cephalodella*, especially *C. catellina* that was exclusively found in the arid region. *Cephalodella catellina* is an eurythermal species reported from both cold and warm regions (Jersabekand & Leitner, 2013; El-Otify & Iskaros, 2015; 2018). Thus, this species appears to be a generalist regarding temperature. To date, the species has only been recorded in Nile area of Egypt (El-Otify & Iskaros, 2015; 2018). A plausible scenario is that resting eggs of this species arrived in the desert ponds from the Nile population and found suitable conditions for maintaining a sustainable population. A genetic characterization of the two populations could help elucidate the genetic distance between them, and allow us to draw secure conclusions about the origin of the Israeli population. From all species, only *Rhinoglena ovigera* was exclusively found in Mediterranean mountain region, marking the first record of this oviparous species of *Rhinoglena* outside of Nearctics (Segers & Walsh, 2017). *Rhinoglena ovigera* was first described in 2017 and has been reported from habitats resembling those of the Mediterranean mountain zone, such as high-altitude ponds exposed to rainfall in the southern USA (Segers & Walsh, 2017).

While most species were widespread across the Mediterranean coast and mountain regions, species of the non-planktonic genus *Lecane* were exclusively found in the Mediterranean coast region. Non-planktonic rotifers, in contrast to planktonic rotifers, deposit their diapausing eggs on a substrate (Serrania-Soto et al., 2011). The extended hydroperiod in this region likely fostered abundant macrophytes and algae, possibly introduced by mobile linkers visiting the ponds to drink, (Parry et al., 2023). These macrophytes/algae provide diverse microhabitats for *Lecane* females to live and attach their eggs. *Epiphanes brachionus*, a recognized indicator species for North America playa habitats (i.e., temporary and semi-permanent water bodies of various sizes; Sublette & Sublette, 1967; Brown et al., 2020), which are comparable in their environmental characteristics (i.e., hydroperiod) to our study system, was found to be extensively distributed in the Mediterranean coast and mountain regions.

Eight pairs of species exhibited positive co-occurrence patterns, suggesting that these species may share similar traits related to habitat selection. The invertebrate community in temporal ponds undergo changes during the wet period (Boix et al., 2016; Hirshberg & Ben-Ami, 2019). In our study, we investigated dormant communities, precluding the investigation of temporal appearance of different species. Thus, it is possible that co-occurring species might have occupied the ponds during different seasons. Two pairs of species displayed a negative co-occurrence, suggesting differences in their ecological traits. Alternatively, successful dispersal might not have occurred to all regions/ponds, or it might have been hindered by short hydroperiods or timing of arrival combined with priority effects (De Meester et al., 2002), as successful dispersal does not necessarily equate to successful colonization (Louette & De Meester, 2004).

Altitude, hydroperiod (i.e., permanency), and mean summer daily maximum temperature significantly affected species richness, diversity, and community composition by. Notably, species richness exhibited a negative correlation with altitude. Previous studies have demonstrated that high altitudes can reduce dissolved oxygen levels, impacting invertebrate communities (Jacobsen et al., 2003). Altitude also negatively affects macrophytes species richness (Rolon & Maltchik, 2006), which serve as microhabitats for certain rotifer species (Duggan et al., 2001) and as secondary dispersal agents (Parry et al., 2023). Furthermore, the species composition of aquatic insects, some of which are predators of rotifers (Hampton & Gilbert, 2001), may also been influenced by altitude (Godon et al., 2017). These findings suggest that altitude may directly or indirectly play a significant role in shaping rotifer communities by affecting macrophytes communities or potential predators.

Species richness and species diversity were further negatively impacted by mean summer daily maximum temperature. This finding is in line with a previous study by Hayee et al. (2015), who demonstrated that rotifers species richness in active communities is negatively affected by air temperature. Increased temperatures coupled with insufficient precipitation can decrease hydroperiod length (Matthews, 2010; Montrone et al., 2019), ultimately leading to a reduced permanency of the habitat patch. Our study revealed that the rotifer community composition is affected by hydroperiod, consistent with former studies on rotifers and other invertebrate communities (Tavernini, 2008; Walsh et al., 2014). During the wet period, the species composition of invertebrates changes greatly, with different species appearing in different times of the season (Boix et al., 2016). Longer hydroperiods have been previously associated with higher species richness of rotifers and crustaceans (Walsh et al., 2014). Therefore, a possible explanation might be that longer hydroperiods allow more time for species to arrive and successfully colonize water bodies, thereby increasing the number of species they accommodate. Conversely, shorter hydroperiod, leading toa longer dry period, not only affect the species assemblage but may also impact the hatching of resting eggs of different rotifers species (Chittapun et al., 2005).

Comparing the three climatic regions, we observed the highest species richness and diversity in the Mediterranean coast region. The ponds in this area are relatively large (375-200000m^2^), and thus often yield a longer hydroperiod. Size has been often correlated with habitat heterogeneity (Tews et al., 2004), potentially facilitating the establishment of diverse and species rich communities. In contrast, ponds located in the arid region are relatively small (0.04-800m^2^), have short hydroperiods, and are often less heterogeneous. They are situated at high altitudes and experience high summer maximum temperatures, making the environment harsher and more unpredictable. Small and unpredictable ponds have been shown to be inhabited dominantly by rotifers, which exhibit an accelerated production of resting eggs (Schröder et al., 2007). Thus, the unpredictability of these ponds may have led to the selection of species that produce resting eggs quickly upon rehydration. These eggs can survive high temperatures, resulting in low species richness and diversity. Additionally, the small size and short hydroperiods in arid regions may have limited the time period for some rotifers species to establish their populations and, thus, lead to a competitive advantage (i.e., priority effects/monopolization) of species that arrived first (De Meester et al., 2002, 2016).

In our exploration of rotifer metacommunity dynamics, we utilized variance partitioning to delve into the underlying factors. Our findings revealed that neither the spatial configuration (2 %) nor the environment (1 %) independently explained a sufficient portion of the rotifer metacommunity variance. When considering both environmental factors (mean summer daily maximum temperature, altitude, and permanency (s/e/p)) and spatial configuration (measured as PCNMs), this proportion slightly increased to 5%.However, the Mantel test revealed a weak but positive correlation between community dissimilarities and geographic distance (i.e., R = 0.119), and environmental heterogeneity (i.e., R = 0.161), with a slightly stronger correlation with environmental heterogeneity (*Supplementary material Figure S1, S2*). This could be partly explained by the linear set-up of the sampling locations (*cf. Figure 1*). Former studies suggest that rotifers, due to their relatively small size, are ubiquitously distributed over large geographic scales (Fontaneto, 2019) by abiotic factors such as wind (Rivas et al., 2019) or biotic vectors (i.e., mobile links), including migratory birds (Green et al., 2023, Moreno et al., 2019, Frisch et al., 2007). Despite Israel being a crucial resting stop for many birds (Lublin et al., 2022), particularly in the northern Mediterranean mountain area (i.e., Hula Lake Park; Gophen, 2023), which may transport organisms/propagules (Green et al., 2023), this study found a stronger effect between community composition and geographic distance (i.e., dispersal limitation) than environmental parameters (i.e., environmental filtering/niche selection). However, since this study was conducted on dormant communities that represent a spatio-temporal accumulation of species, we cannot derive direct conclusions of community processes, such as environmental filtering/niche selection, species interactions, dispersal limitation and ecological drift. Future comparative surveys on active communities in the water column will be necessary to validate these findings. Nonetheless, our findings suggest that dispersal limitation over an area of 260×60 km combined with environmental filtering/niche selection, appear to be the dominant processes in rotifer community assemblage in Israel. This is consistent with studies conducted on small freshwater bodies in Europe, North and Central America (e.g., Gálvez et al., 2020, Florencio et al., 2014, Gilbert et al., 2023). The relatively large amount of unexplained variance (92 %) likely stems from the fact that only dormant communities were sampled. These dormant communities represent a spatio-temporal accumulation that could mask the underlying community processes, as they typically occur during the re-wetting phase of a pond.

## Conclusions and Limitations

In our study, we aimed to estimate the diversity of monogonont rotifers in Israel by rehydrating sediment samples from 30 temporary ponds. This approach allowed us to conduct a cost-effective assessment of rotifer community diversity, but we acknowledge that our use of a fixed temperature of 20°C for sediment rehydration may have introduced some bias against species that have other temperature hatching optima. While we identified altitude, hydroperiod, and mean summer daily maximum temperature as key environmental drivers of rotifer community composition and species richness in Israel, our study only examined dormant communities, and thus further research on a spatio-temporal scale of the active communities in the water column is necessary to validate our findings and strengthen the ecological factors behind. Despite these limitations, we were able to discover 25 previously unknown rotifer species in the area, highlighting the importance of this approach for assessing rotifer community diversity in Israel. To gain a more comprehensive understanding of the country’s rotifer fauna, future research should expand to include other freshwater habitats such as lakes, permanent ponds, ditches, puddles, and wetlands.

## Author contribution statement

Conceptualization: OH, SP, FB-A. Developing methods: OH, SP, KK, FB-A. Data analysis: OH, SP, KK. Preparation of figures and tables: OH, SP, KK. Conducting the research: OH, SP. Data interpretation and writing: OH, SP, KK, FB-A.

## Acknowledgments

We are grateful to Henrik Segers for his help in identifying *R. ovigera*.

## Data Availability Statement

Data supporting this study as well as scripts used for the analyses are available on figshare, DOI: 10.6084/m9.figshare.24347584.

