## supplementary for "The gems of the Middle-East: Unveiling the biodiversity of Monogonont rotifers in temporary waterbodies of Israel": Supplementary_material_13.7.23.docx

Table S1: Summary of environmental parameters of the sampled temporary ponds. Listed are sampling region, sampling location, pond size [m^2^], altitude [m], annual precipitation [mm^2^], daily evaporation [mm^2^], permanency ratios (e/p; s/e/p), wind velocity [m/s], number of neighboring ponds (5 km radius and 15 km radius).

| **Pond number** | **longitude** | **latitude** | **Region** | **Size (m^2)** | **Altitude (m)** | **Mean summer daily maximum temperature (C)** | **Mean winter daily minimum temperature (C)** | **Annual precipitation (mm^2)** | **Daily evaporation (mm^2)** | **Annual evaporation (mm^2)** | **Ratio e/p** | **Ratio size/(e/p)** | **Wind velocity (m/s)** | **Neighbours 5km radius** | **Neighbours 15km radius** |
| --- | --- | --- | --- | --- | --- | --- | --- | --- | --- | --- | --- | --- | --- | --- | --- |
| 1 | 33.04938 | 35.48955 | Mediterranean mountain | 6540 | 681 | 29.89 | 5.43 | 149 | 9 | 3285 | 22 | 297.32 | 1 | 5 | 7 |
| 2 | 33.02969 | 35.49055 | Mediterranean mountain | 4900 | 722 | 33.02 | 6.86 | 102 | 9 | 3285 | 32.34 | 151.52 | 1 | 5 | 7 |
| 3 | 33.017 | 35.49 | Mediterranean mountain | 6400 | 809 | 36.14 | 8.28 | 125 | 9 | 3285 | 26.21 | 244.14 | 1 | 5 | 7 |
| 4 | 32.41104 | 34.8981 | Mediterranean coast | 200000 | 5 | 31.6 | 8.62 | 109 | 7.36 | 2686.4 | 24.71 | 8092.61 | 3.9 | 2 | 5 |
| 5 | 32.41104 | 34.8981 | Mediterranean coast | 8000 | 5 | 31.6 | 8.62 | 109 | 7.36 | 2686.4 | 24.71 | 323.7 | 3.9 | 2 | 5 |
| 6 | 32.25921 | 34.84769 | Mediterranean coast | 20000 | 12 | 31.22 | 9.79 | 105 | 7.36 | 2686.4 | 25.69 | 778.37 | 3.9 | 4 | 8 |
| 7 | 32.25921 | 34.84769 | Mediterranean coast | 10000 | 12 | 31.22 | 9.79 | 105 | 7.36 | 2686.4 | 25.69 | 389.18 | 3.9 | 4 | 8 |
| 8 | 32.2439 | 34.8545 | Mediterranean coast | 29400 | 30 | 31.22 | 9.79 | 105 | 7.36 | 2686.4 | 25.69 | 1144.2 | 3.9 | 4 | 9 |
| 9 | 32.231 | 34.827 | Mediterranean coast | 6300 | 25 | 30.85 | 11.69 | 101 | 7.36 | 2686.4 | 26.49 | 237.84 | 3.6 | 4 | 9 |
| 10 | 32.17067 | 34.82421 | Mediterranean coast | 776 | 24 | 30.85 | 11.69 | 101 | 7.36 | 2686.4 | 26.49 | 29.3 | 3.2 | 1 | 8 |
| 11 | 32.17067 | 34.82421 | Mediterranean coast | 345 | 24 | 30.85 | 11.69 | 101 | 7.36 | 2686.4 | 26.49 | 13.02 | 3.2 | 1 | 8 |
| 12 | 32.13583 | 34.79444 | Mediterranean coast | 3000 | 22 | 30.74 | 12.1 | 76 | 7.4 | 2701 | 35.49 | 84.52 | 3.2 | 2 | 10 |
| 13 | 32.03514 | 34.77243 | Mediterranean coast | 572 | 0 | 30.74 | 12.1 | 76 | 7.4 | 2701 | 35.49 | 16.12 | 2.6 | 1 | 2 |
| 14 | 32.03514 | 34.77243 | Mediterranean coast | 429 | 0 | 30.74 | 12.1 | 76 | 7.4 | 2701 | 35.49 | 12.09 | 2.6 | 1 | 2 |
| 15 | 31.91861 | 34.73944 | Mediterranean coast | 37400 | 0 | 31.46 | 9.69 | 113 | 7.4 | 2701 | 23.97 | 1560.11 | 2.6 | 0 | 3 |
| 16 | 31.7251 | 34.6761 | Mediterranean coast | 4200 | 40 | 32.9 | 8.48 | 96 | 8.71 | 3179.15 | 33.23 | 126.39 | 1.6 | 3 | 4 |
| 17 | 31.7251 | 34.6761 | Mediterranean coast | 5000 | 40 | 32.9 | 8.48 | 96 | 8.71 | 3179.15 | 33.23 | 150.46 | 1.6 | 3 | 4 |
| 18 | 31.7012 | 34.71416 | Mediterranean coast | 10000 | 54 | 32.54 | 9.27 | 85 | 8.71 | 3179.15 | 37.54 | 266.36 | 1.6 | 2 | 6 |
| 19 | 31.06608 | 35.26943 | Arid | 0.0777 | 256 | 37.82 | 11.415 | 11 | 7.09571429 | 2589.935714 | 227.05 | 0 | 3.9 | 2 | 2 |
| 20 | 31.06493 | 35.2714 | Arid | 0.45041 | 229 | 37.82 | 11.415 | 11 | 7.09571429 | 2589.935714 | 227.05 | 0 | 3.9 | 2 | 2 |
| 21 | 31.06596 | 35.27493 | Arid | 0.0422 | 200 | 37.82 | 11.415 | 11 | 7.09571429 | 2589.935714 | 227.05 | 0 | 3.9 | 2 | 2 |
| 22 | 30.95222 | 35.0253 | Arid | 720 | 508 | 33.03 | 6.72 | 12 | 11.09 | 4047.85 | 329.09 | 2.19 | 3.4 | 0 | 0 |
| 23 | 30.66453 | 34.81074 | Arid | 18 | 796 | 33.03 | 6.72 | 12 | 11.09 | 4047.85 | 329.09 | 0.05 | 3 | 0 | 2 |
| 24 | 30.61941 | 34.89112 | Arid | 800 | 478 | 33.03 | 6.72 | 12 | 11.09 | 4047.85 | 329.09 | 2.43 | 3 | 0 | 1 |
| 25 | 30.59351 | 34.70911 | Arid | 225 | 809 | 33.03 | 6.72 | 12 | 11.09 | 4047.85 | 329.09 | 0.68 | 3 | 0 | 3 |
| 26 | 30.526 | 34.6119 | Arid | 2.25 | 975 | 33.03 | 6.72 | 12 | 11.09 | 4047.85 | 329.09 | 0.01 | 3 | 1 | 2 |

Table S2: Permanency and altitude categories which were used for some summary statistics (i.e., NMDS, ANOVA).

| **Parameter** | **Values** | **Category** |
| --- | --- | --- |
| Permanency  (s/e/p) | 0-10 | 1 |
|  | 11-100 | 2 |
|  | 101-200 | 3 |
|  | 201-1000 | 4 |
|  | >1000 | 5 |
| Altitude | 1-100m | 1 |
|  | 101-400m | 2 |
|  | 401-600m | 3 |
|  | 601-800m | 4 |
|  | >800 | 5 |

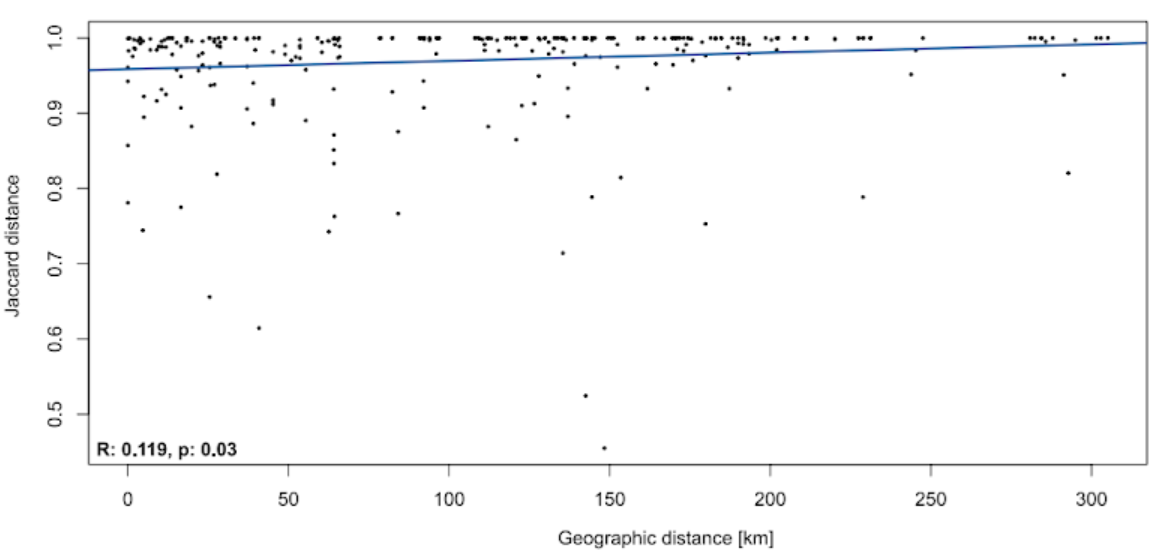

Figure S1: Correlogram of geographic distance and Jaccard dissimilarity of the hatched communities of the studied temporary ponds (n = 26). The line indicates the relation between geographic distance [km] and Jaccard distances.

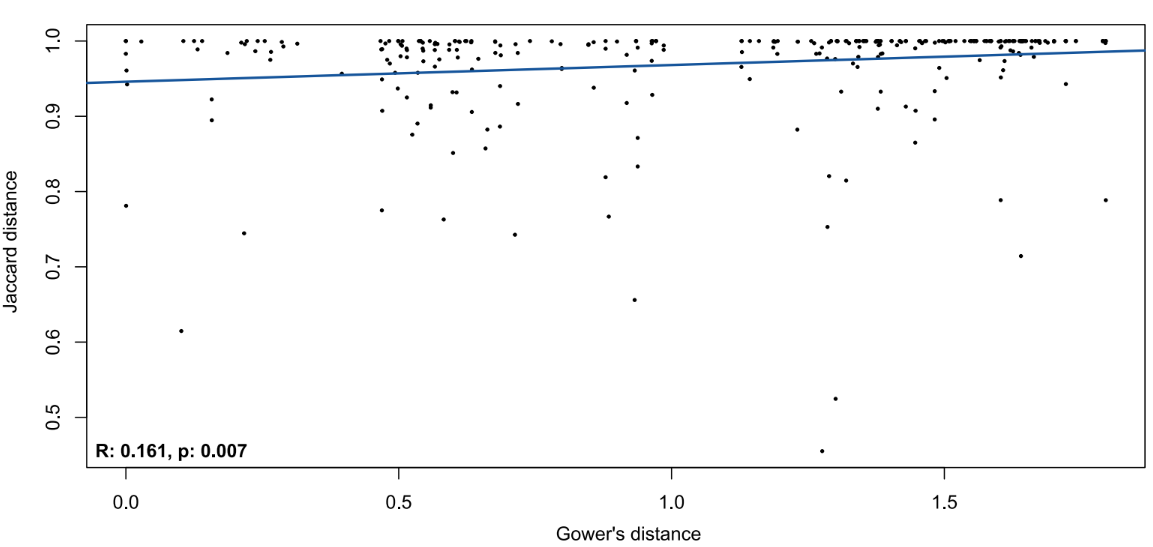

Figure 2S: Correlogram of environmental distance (i.e., Gower’s distance) and Jaccard dissimilarity of the hatched communities of the studied temporary ponds (n = 26). The line indicates the relation between Gower distance and Jaccard distances.
